# BECLIN-1 is Essential for the Maintenance of Gastrointestinal Epithelial Integrity by Regulating Endocytic Trafficking, F-actin Organization and Lysosomal Function

**DOI:** 10.1101/2024.11.28.625769

**Authors:** Juliani Juliani, Sharon Tran, Tiffany J. Harris, Peter De Cruz, Sarah L. Ellis, Paul A. Gleeson, John M. Mariadason, Kinga Duszyc, Alpha S. Yap, Erinna F. Lee, Walter D. Fairlie

**Author notes:** **Co-corresponding authors**, Erinna Lee, Walter Fairlie.

## Abstract

Disrupted intestinal homeostasis and barrier function are key contributors to the development of various diseases, including inflammatory bowel disease (IBD). BECLIN-1, a core component of two class III phosphatidylinositol 3-kinase (PtdIns3K) complexes, has a dual role in autophagy and endocytic trafficking. Emerging evidence suggests it is involved in maintaining intestinal integrity which involves its endocytic trafficking function. To gain insights into the fatal gastrointestinal (GI) phenotype observed in BECLIN-1 knockout adult mice, organoids derived from these animals were used to investigate the role of BECLIN-1 in GI epithelial function. BECLIN-1 deletion led to disrupted localization of CADHERIN-1/E-CADHERIN to adherens junctions (AJs) and OCCLUDIN to tight junctions (TJs), resulting in the mislocalization of these critical barrier proteins. Lysosomal dysfunction was also observed, characterized by impaired cargo degradation. In addition, the filamentous actin (F-actin) cytoskeleton became disorganized though this was not associated with changes in the interaction between F-actin and CATENIN BETA-1/BETA-CATENIN nor in alterations in BETA-CATENIN localization, though BETA-CATENIN levels were reduced. These effects were all less pronounced or absent in organoids lacking the autophagy-only regulator ATG7 (autophagy related 7), further emphasizing the involvement of BECLIN-1’s protein trafficking role in the maintenance of gut homeostasis and barrier function. These findings provide new insights into processes involved in epithelial dysfunction, deepening our understanding of mechanisms underlying intestinal disease.

## Introduction

The maintenance of intestinal homeostasis relies on a complex interplay of cellular processes that provide a protective barrier and ensure effective nutrient absorption and immune cell regulation [1, 2]. Over the past two decades, research into the role of autophagy in gut health has grown significantly, stemming from landmark discoveries that uncovered polymorphisms in the core autophagy gene *ATG16L1* (autophagy related 16 like 1) linked to Crohn’s disease [3–6]. This breakthrough, together with subsequent mechanistic studies, provided a deeper understanding of the genetic basis of inflammatory bowel disease (IBD) and highlighted the critical role autophagy plays in intestinal health [7, 8]. Subsequently, a multitude of genetic models in which key autophagy regulators were modified have been generated to decipher the role of autophagy in intestinal epithelial cells (IECs) [1]. Specifically, these studies have elucidated the contribution of autophagy to immune responses in the gut [9, 10], intestinal barrier integrity [11–14], gut microbiota biology [15–17] as well as the pathogenesis of intestinal diseases [18, 19] and therapeutic potential for treating gut-related disorders [13, 20, 21].

BECLIN-1 is a core component of two class III PtdIns3K complexes and acts as a signal integration hub with roles in both autophagy and endocytic trafficking [22]. Accordingly, BECLIN-1 has been implicated in the pathogenesis of numerous diseases, including various cancers (breast, ovarian, prostate, colorectal), Alzheimer’s disease, cystic fibrosis, sepsis and others [22–30]. Germline deletion of BECLIN-1 results in an embryonic lethal phenotype in contrast to constitutive deletion of autophagy conjugation proteins such as ATG5 (autophagy related 5) or ATG7 which results in neonatal lethality [24, 31]. The disparity suggests an essential role for BECLIN-1 beyond autophagy during development.

In a recent study, we circumvented the embryonic lethality of germline BECLIN-1 deletion using a conditional BECLIN-1 knockout mouse model, which enabled inducible, whole body and intestinal epithelial cell-specific BECLIN-1 deletion in adult mice. This led to a fatal enteritis phenotype that we attributed, using intestinal organoids, to defects in endocytic trafficking that resulted in mislocalization of the critical junctional protein, E-CADHERIN. This severe phenotype was not observed in equivalent studies involving deletion of the autophagy regulator ATG7, which is not associated with trafficking, underscoring a unique and essential function for BECLIN-1 in maintaining intestinal homeostasis. [11, 32]. Additional published reports have also implicated BECLIN-1 in maintaining intestinal homeostasis though these primarily focused on the colon and relied on autophagy-proficient experimental systems such as Tat-Beclin1 peptide treatment and *Becn1*^F121A^ mice [13, 17].

In this study, we show that the role of BECLIN-1 in maintaining small intestinal integrity extends beyond the localization of E-CADHERIN at the adherens junctions (AJs) and is also critical in forming tight junctions (TJs). In addition, we show that its deletion leads to lysosomal dysfunction and cytoskeletal disruption, further compromising the epithelial barrier. Combined, these data provide a comprehensive understanding of how BECLIN-1 supports intestinal homeostasis by coordinating junctional integrity, endocytic trafficking, and cellular architecture to preserve the epithelial barrier.

## Results

### BECLIN-1 loss leads to mislocalization of OCCLUDIN that mirrors E-CADHERIN mislocalization in intestinal organoids

Our published studies showed that BECLIN-1 deletion in intestinal organoids leads to the mislocalization of E-CADHERIN at the AJ, which we hypothesized contributes to increased intestinal epithelial permeability in mice [11]. However, in addition to AJs, epithelial barrier integrity is maintained by the coordinated functions of several junctional complexes, including TJs and desmosomes [2]. Interestingly, a previous study demonstrated that whilst BECLIN-1 knockdown disrupted OCCLUDIN trafficking, it unexpectedly reduced TJ permeability in Caco-2 colon cancer cells grown in 2D cultures [13]. Given that our studies had suggested that BECLIN-1 loss compromises the epithelial barrier, we utilized our more physiologically relevant 3D small intestine-derived organoids, to first investigate the role of BECLIN-1 in OCCLUDIN localization at TJs. These experiments used organoids at an earlier developmental stage (Day 5) compared to our previous work, allowing us to focus on events preceding overt BECLIN-1 deletion-induced organoid death. For clarity, references to the apical plasma membrane refer to the apicolateral junction, while the lateral membrane extends downwards, stopping just before the basolateral junction (see Supplemental Figure S1A for details).

As we showed previously [11], BECLIN-1 deletion led to a significant reduction in apicolateral E-CADHERIN localization in organoids derived from intestinal epithelial-specific BECLIN-1-deficient (*Becn1^ΔIEC^*) mice compared to organoids derived from their wild-type littermates (*Becn1^wtIEC^*) (Figure 1A, B). This was accompanied by increased E-CADHERIN localization along the lateral membranes (Figure 1A, C) and enhanced cytoplasmic accumulation of E-CADHERIN (Figure 1A, D). OCCLUDIN, which is typically localized to the apicolateral region above E-CADHERIN [13, 33], displayed a similar mislocalization pattern in *Becn1^ΔIEC^* organoids. This included a significant loss of apicolateral OCCLUDIN (Figure 1A, E), with increased lateral (Figure 1A, F) and cytoplasmic accumulation (Figure 1A, G), mirroring the changes seen with E-CADHERIN.

**Figure 1.**
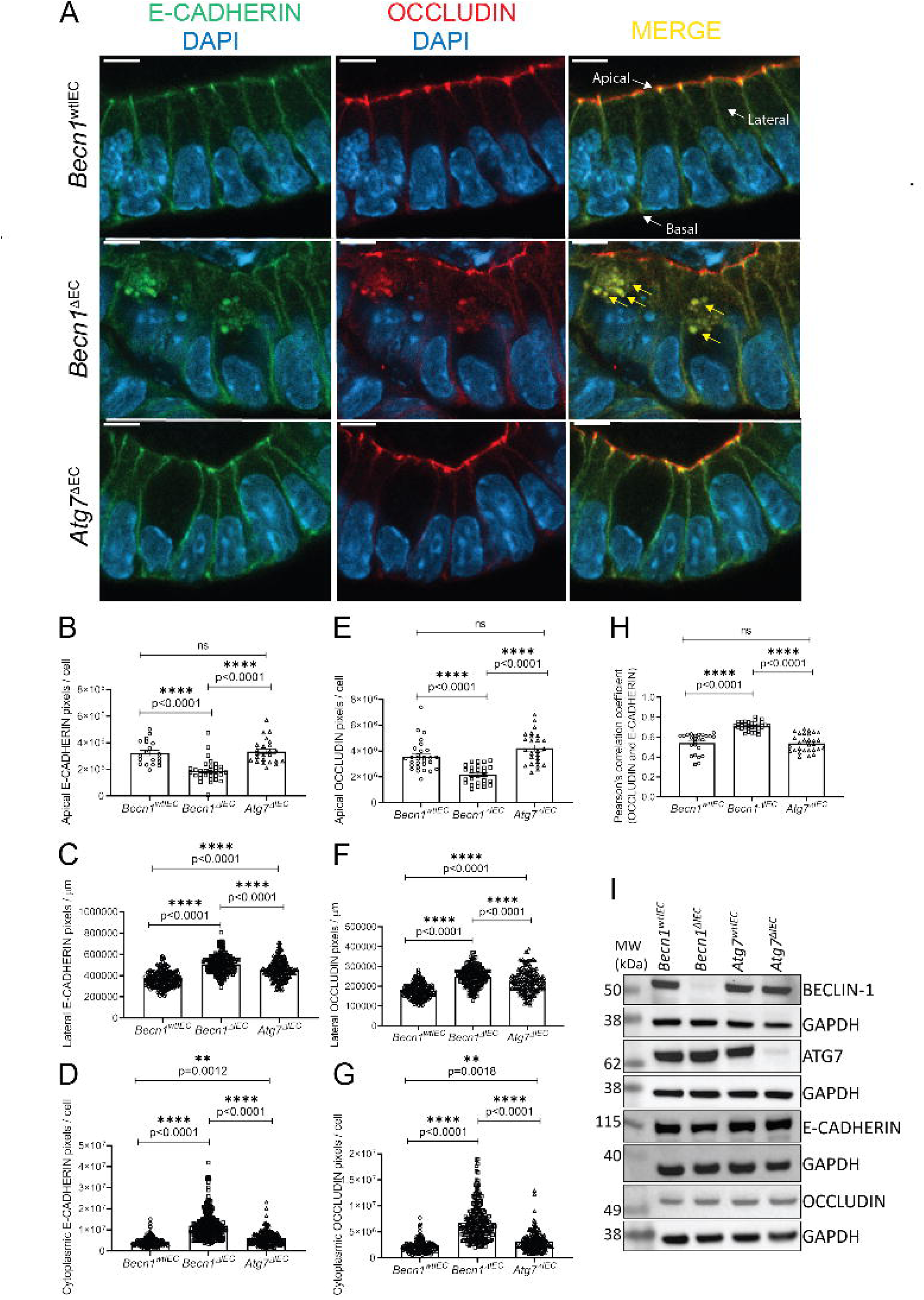
Mislocalization of E-CADHERIN and OCCLUDIN following BECLIN-1 loss in intestinal organoids. (**A**) Representative fields from whole-mount immunofluorescent staining of Day 5 post-4-HT-treated *Becn1^wtIEC^*, *Becn1^ΔIEC^*, and *Atg7^ΔIEC^* intestinal organoids. The absence of BECLIN-1, but not ATG7, leads to the mislocalization of OCCLUDIN, mirroring that of E-CADHERIN, as detected by immunofluorescence. The mislocalized junctional proteins appear as aberrantly enlarged cytoplasmic puncta, which colocalize with one another. Quantification of immunofluorescence staining shows. Yellow arrows indicate representative enlarged cytoplasmic puncta. Labelled white arrows provide orientation of key membranes for all subsequent figures. (**B**) BECLIN-1 loss results in a significant reduction in apical membrane localization of E-CADHERIN with concurrent accumulation in (**C**) lateral membrane and (**D**) cytoplasmic membranes. These changes are mirrored in the (**E**) apical, (**F)** lateral, and (**G**) cytoplasmic localization of OCCLUDIN. (**H**) The loss of BECLIN-1, but not ATG7, also results in increased co-localization between E-CADHERIN and OCCLUDIN as measured using Pearson’s colocalization coefficient. (**I**) There was no change in total E-CADHERIN and OCCLUDIN protein levels between *Becn1^ΔIEC^*, *Atg7^ΔIEC^* and their wild-type counterparts, as determined by Western blotting. GAPDH (glyceraldehyde-3-phosphate dehydrogenase) was used as a loading control for the blot directly above each GAPDH panel. Molecular weight (MW) markers indicate the relative size of the detected protein. Data are representative of at least *n = 3* biological replicates. Graphs show the mean ± S.E.M. Statistical significance was determined by ordinary one-way ANOVA for all comparisons. Scale bar = 10 µm.

Interestingly, deletion of ATG7 in intestinal organoids also caused mislocalization of both E-CADHERIN and OCCLUDIN, though these changes were less pronounced than those observed in *Becn1^ΔIEC^*organoids (Figure 1A-G). In *Atg7^ΔIEC^* organoids, the lateral and cytoplasmic accumulation of both proteins increased (Figure 1A, C, D, F, G), but apicolateral localization remained largely intact (Figure 1A, B, E), indicating that the significantly impaired apicolateral localization of these junctional proteins is unique to BECLIN-1 deletion.

Co-localization analysis further revealed a significantly higher overlap of E-CADHERIN and OCCLUDIN in *Becn1^ΔIEC^* organoids (i.e. higher Pearson’s colocalization coefficient) compared to both *Becn1^wtIEC^*and *Atg7^ΔIEC^* organoids, suggesting abnormal accumulation of these proteins, most likely within defective RAB5A^+ve^ (Ras-related protein Rab-5a) early endosomes, as we previously reported [11] (Figure 1A, H). Importantly, no significant differences in the total cellular levels of E-CADHERIN and OCCLUDIN were observed across the genotypes (Figure 1I), suggesting that the observed mislocalization reflects trafficking defects rather than changes in expression levels.

In conclusion, the loss of BECLIN-1-mediated endocytic trafficking disrupts not only E-CADHERIN localization but also leads to similar mislocalization of OCCLUDIN. While ATG7 deletion also causes some mislocalization, it is less severe compared to BECLIN-1 loss, particularly with respect to apicolateral localization. These findings provide further insights into our previous observation where increased intestinal permeability was seen in *Becn1^wtIEC^* but not *Atg7^ΔIEC^*mice [11].

### BECLIN-1 deletion more profoundly impairs lysosomal function compared to ATG7 deletion in organoids

We previously demonstrated that BECLIN-1 loss impairs both autophagy flux and disrupts early and late endosome activity [11]. To extend our understanding of these defects in the intestinal epithelium, we next investigated whether BECLIN-1 deletion also affects the lysosomal compartment, the final destination for many cargoes delivered through the endocytic and autophagy pathways. Lysosomal dysfunction could provide a mechanistic link between the observed mislocalization of junctional proteins and defective degradation pathways, contributing to the cellular stress seen in BECLIN-1-deficient organoids [11]. Here, we used LysoTracker Red^TM^ DND-99 to label acidic organelles and Dequenched Green Bovine Serum Albumin (DQ^TM^ Green BSA), a fluorescent cargo that becomes activated upon degradation in lysosomes. In the subsequent sections, LysoTracker Red^TM^ DND-99 and DQ^TM^ Green BSA will be referred to as LysoTracker Red and DQ-BSA, respectively.

At day 5 post-gene deletion, lysosome numbers measured by LysoTracker Red puncta were unchanged across *Becn1^wtIEC^*, *Becn1^ΔIEC^*, and *Atg7^ΔIEC^* organoids (Figure 2A, B). However, *Becn1^ΔIEC^*organoids showed a significant reduction in DQ-BSA particles, indicating impaired cargo degradation (Figure 2A, C). A similar trend (*p =* 0.0942) was observed in *Atg7^ΔIEC^*organoids, though to a lesser extent (Figure 2A, C). This suggests that while BECLIN-1 and ATG7 deletion does not impact the overall lysosome number, cargo (DQ-BSA) delivery to lysosomes or lysosomal proteolytic activity is impaired. This is confirmed by the significant reduction in Mander’s M1 overlap coefficient (LysoTracker Red overlapping DQ-BSA fluorophores) in both *Becn1^ΔIEC^*and *Atg7^ΔIEC^* organoids compared to *Becn1^wtIEC^*organoids (Figure 2A, D).

**Figure 2.**
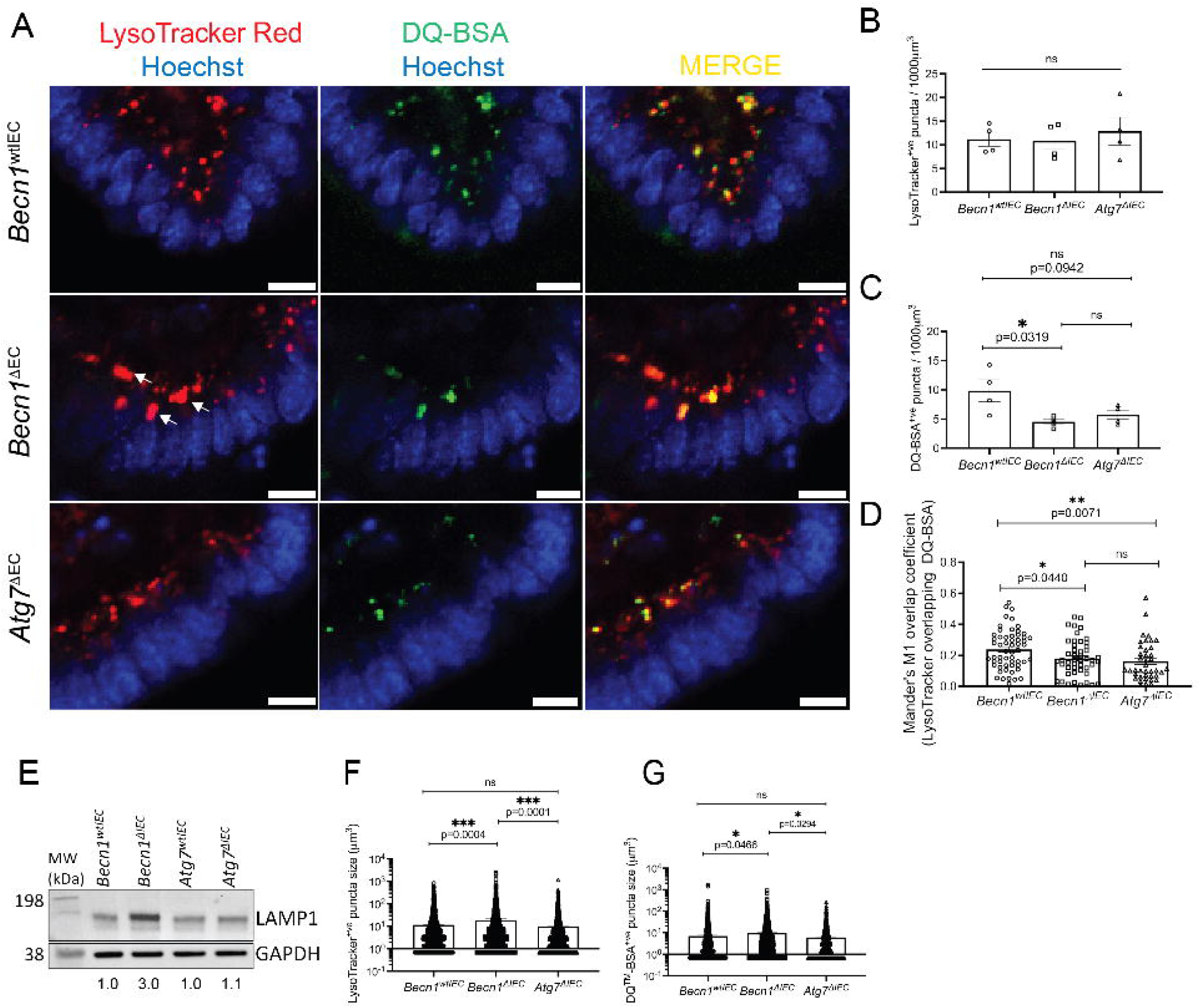
BECLIN-1, but not ATG7, loss led to enlarged lysosomes. **A)** Representative fields from live imaging of intestinal organoids at Day 5 post-4-HT treatment using Lysotracker Red (red, marking lysosomes) and DQ-BSA (green, a marker of cargo degradation). Arrows indicate examples of enlarged lysosomes. Quantification of these data revealed (**B**) no significant change in lysosome numbers, but (**C**) a reduction in cargo degradation activity following BECLIN-1 loss. Additionally, there was (**D**) decreased colocalization between Lysotracker Red and DQ-BSA, quantified using Mander’s overlap coefficient. Colocalization analysis was performed across all z-sections per organoid. (**E**) BECLIN-1 loss, but not ATG7 loss, led to increased LAMP1 levels, as detected *via* Western blotting. GAPDH was used as a loading control. Numbers below each lane represent levels of LAMP1 relative to the corresponding wild-type organoids for either *Becn1* or *Atg7*-deficient organoids, as determined by densitometry analysis. Values are the averages of *n*= 4 replicate experiments. Molecular weight (MW) markers indicate the relative size of the detected protein. There was also (**F**) an increase in Lysotracker Red and **G)** DQ-BSA puncta size (quantified from >1900 puncta). Data were representative of *n =* 4 biological replicates. Graphs show the mean ± S.E.M. Significance was determined by ordinary one-way ANOVA for all comparisons. Scale bar = 10 µm.

Despite these shared defects between the BECLIN-1 and ATG7-deleted organoids, distinct differences were apparent. Specifically, *Becn1^ΔIEC^*organoids exhibited significantly enlarged lysosomes (i.e. LysoTracker Red^+ve^ puncta), accompanied by 3-fold increased levels of the lysosomal membrane protein, LAMP1 (lysosomal-associated membrane protein 1), levels, unlike *Atg7^ΔIEC^* organoids (Figure 2A, E, F). This enlargement was also accompanied by larger DQ^-^BSA (though fewer) puncta (Figure 2A, C, G), suggesting that although some cargo reaches the lysosome, it is not efficiently degraded, leading to the accumulation of the DQ-BSA^+ve^ fluorescent signal.

In summary, BECLIN-1 loss leads to defective lysosomal cargo degradation and enlargement of lysosomes, whereas ATG7 deletion causes milder lysosomal defects. This highlights a unique role for BECLIN1 versus ATG7 in maintaining lysosomal function.

### Dynamic assessment of endocytic trafficking in BECLIN-1 and ATG7-Deleted Mouse Embryonic Fibroblasts

Static imaging of RAB markers provides insights into endosomal distribution, but real-time analysis is crucial for understanding the dynamic nature of endocytic trafficking. To monitor cargo movement, live uptake assays were conducted using the pH-sensitive dye (pHrodo) conjugatated to Dextran (bulk fluid-phase endocytosis) or Transferrin (receptor-mediated endocytosis), as well as AlexaFluor^TM^ 594 -conjugated wheat germ agglutinin (WGA) (clathrin-dependent and independent pathways). Despite multiple attempts using different protocols, these fluorescent probes did not penetrate the 3D matrix in our organoid system. Therefore, to provide a system more amenable to these experiments, we utilized SV40 large T antigen-transformed mouse embryonic fibroblasts (MEFs) derived from BECLIN-1 and ATG7 knockout mouse embryos. Tamoxifen metabolite, 4-hydroxytamoxifen (4-HT) treatment resulted in successful deletion of BECLIN-1 (*Becn1^-/-^*) and ATG7 (*Atg7^-/-^*) at day 5 post-treatment (Supplemental Figure S2A). Similar to that observed in mouse IECs [11], *Becn1^-/-^* and *Atg7^-/-^* MEFs demonstrated defective basal autophagy flux, characterized by a pronounced accumulation of total SQSTM1/P62 (sequestosome 1) and MAP1LC3B (microtubule-associated protein 1 light chain 3 beta)-I and -II levels or a reduced MAP1LC3B-II:MAP1LC3B-I ratio compared to respective wild-type controls (Supplemental Figure S2B).

Real-time imaging of *Becn1^-/-^* MEFs showed there was a significant reduction in Dextran and WGA-positive puncta within the cytoplasm compared to wild-type controls and *Atg7^-/-^* MEFs (Supplemental Figure S2C-E, H). Instead, cargo strikingly accumulated near the plasma membrane following internalization, indicating a block in the early trafficking stages. Similarly, transferrin uptake and trafficking *via* a more tightly regulated clathrin-dependent receptor-mediated endocytic process were defective compared to wildtype and *Atg7^-/-^* MEFs, with cargo stalling near the plasma membrane and failing to progress into endosomes (Supplemental Figure S2F, G).

These results suggest that BECLIN-1, unlike ATG7, is critical for the proper progression of cargo through the early endocytic stages. The consistent disruption across different pathways (i.e. fluid-phase, receptor-mediated and clathrin-independent) indicates a potential common point of failure at the level of the RAB5A^+ve^ early endosome, consistent with the effect of BECLIN-1 deletion on this compartment in our recent studies [11].

### Disruption of ACTIN cytoskeleton integrity in intestinal organoids upon BECLIN-1, but not ATG7, deletion

Having established that BECLIN-1 loss disrupts endocytic trafficking and the correct localization of junctional proteins such as E-CADHERIN and OCCLUDIN, we next investigated the F-actin cytoskeleton. F-actin filaments are fundamental to maintaining intestinal epithelial structure and function, including junction assembly, stability, and endocytic trafficking [34]. The cortical actin network, particularly the apical F-actin belt, is directly associated with TJs and AJs, and the correct localisation of TJs and AJs is important for the formation of the F-actin cytoskeleton. Additionally, F-actin filaments play a key role in assembly and sorting of tubular carriers as well as endosome maturation and movement [34]. Given the interconnected roles of the F-actin cytoskeleton in junctional protein localization and trafficking, we hypothesized that BECLIN-1 loss might also affect F-actin organization, contributing to the observed disruption in junctional protein integrity and barrier function.

Wholemount staining of intestinal organoids for F-actin and E-CADHERIN revealed a marked reduction in the apical F-actin belt in *Becn1^ΔIEC^* organoids compared to *Becn1^wtIEC^* and *Atg7^ΔIEC^* organoids (Figure 3A, B). This loss of the F-actin belt was accompanied by decreased co-localization of F-actin with E-CADHERIN at apicolateral junctions, as evidenced by Pearson’s correlation coefficient (Figure 3A, C).

**Figure 3.**
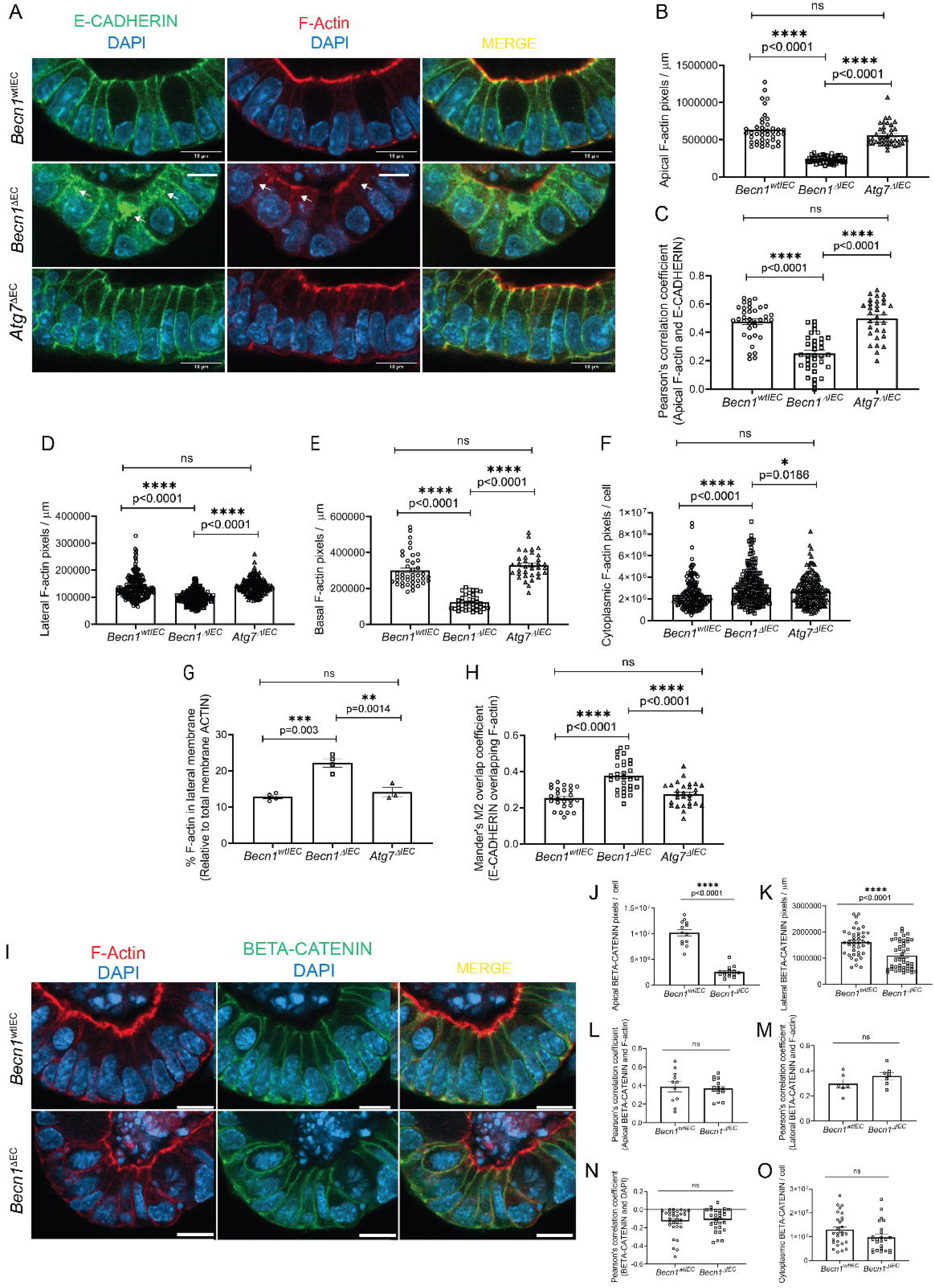
Disruption of the F-actin cytoskeleton following BECLIN-1 loss. (**A**) Representative fields from whole-mount staining of intestinal organoids with E-CADHERIN and F-actin. Arrows indicate regions of mislocalized E-CADHERIN or F-actin. Quantification of these data revealed (**B**) significant depletion of apical F-actin staining and **C)** reduced colocalization of E-CADHERIN and F-actin at the apicolateral junction, as measured by Pearson’s correlation coefficient, following BECLIN-1 loss. Additionally, (**D**) lateral and (**E**) basal F-actin expression in intestinal organoids was significantly decreased, while (**F**) cytoplasmic accumulation of F-actin was evident. (**G**) BECLIN-1 loss led to an increased ratio of lateral F-actin relative to total membrane-associated F-actin and (**H**) increased overlap of E-CADHERIN and F-actin at the lateral membrane and cytoplasm. (**I**) Representative fields from whole-mount immunostaining of F-actin and BETA-CATENIN also revealed a significant decrease in (**J**) apical and (**K**) lateral membrane staining of BETA-CATENIN following BECLIN-1 loss, with (**L, M**) no changes in the colocalization of F-actin and BETA-CATENIN in both regions. (**N**) There were also no significant changes in the colocalization of BETA-CATENIN with DAPI and (**O**) no increase in BETA-CATENIN cytoplasmic accumulation following BECLIN-1 loss. Data were representative of at least *n = 3* biological replicates. Graphs show the mean ± S.E.M. Statistical significance was determined by ordinary one-way ANOVA for B-H and student’s unpaired t-test for J-O. Scale bar = 10 µm.

Further analysis revealed that *Becn1^ΔIEC^* organoids also had a significant reduction in lateral and basal F-actin, associated with increased cytoplasmic accumulation of F-actin compared to controls (Figure 3D-F). However, there was a redistribution of F-actin towards the lateral membrane as indicated by the increased ratio of lateral membrane-associated F-actin relative to total membrane-associated F-actin (Figure 3G). This increased lateral and cytoplasmic F-actin overlapped (Figure 3H) with an increase in lateral and cytoplasmic E-CADHERIN in *Becn1^ΔIEC^* organoids (Figure 1C, D), which was not observed in *Becn1^wtIEC^* and *Atg7^ΔIEC^* organoids.

These findings highlight the critical role of the F-actin cytoskeleton in maintaining epithelial structure and integrity, which becomes severely compromised upon BECLIN-1 loss. The disruption of F-actin cytoskeleton would not only exacerbate epithelial barrier dysfunction, it would contribute to impairment in endocytic trafficking, leading to the destabilisation of cell junctions.

### Loss of F-actin in *Becn1^ΔIEC^* organoids is not attributed to defects in the BETA-CATENIN-F-actin interaction

The recruitment and organization of F-actin at junctional sites are partially regulated by BETA-CATENIN, a key component of the Wnt signalling pathway that interacts with E-CADHERIN [35]. As BETA-CATENIN mediates cytoskeletal remodelling and junction integrity by forming part of the multiprotein complex with CATENIN ALPHA 1 that links E-CADHERIN to F-actin, we investigated whether the loss of F-actin in *Becn1^ΔIEC^*organoids is associated with changes in junctional BETA-CATENIN. Additionally, BETA-CATENIN plays a dual role in Wnt signalling, translocating to the nucleus to regulate gene expression related to the cytoskeletal organization. Hence, we also examined whether BECLIN1 loss affected BETA-CATENIN localization or its nuclear translocation.

Wholemount staining showed a significant reduction in apical and lateral BETA-CATENIN signals in *Becn1^ΔIEC^* organoids compared to controls (Figure 3I-K). This reduction mirrored the loss of F-actin in these regions (Figure 3B, D). However, despite the decrease in BETA-CATENIN and F-actin levels, there was no significant difference in their co-localization between *Becn1^ΔIEC^* and *Becn1^wtIEC^* organoids (Figure 3I, L, M). This suggests that the BETA-CATENIN-F-actin interaction remains intact.

To investigate whether the loss of BETA-CATENIN from the membrane was associated with increased nuclear translocation, we assessed BETA-CATENIN localization in the nucleus. No significant increase in nuclear BETA-CATENIN was observed between *Becn1^ΔIEC^* and *Becn1^wtIEC^*organoids (Figure 3I, N). Additionally, there was no evidence of cytoplasmic accumulation of BETA-CATENIN, suggesting that normal degradation processes involved in BETA-CATENIN turnover of excess protein (i.e. BETA-CATENIN not bound to E-CADHERIN or WNT signalling) are not affected [36] (Figure 3I, O).

Combined, these data indicate that the disruption of the F-actin cytoskeleton is not due to impairments in BETA-CATENIN-mediated pathways, such as its role in linking E-CADHERIN to F-actin or in Wnt signalling. Instead, it suggests that BECLIN-1 influences F-actin organization through alternative mechanisms, highlighting its unique contribution to maintaining epithelial integrity.

## Discussion

The intestinal epithelium forms a critical barrier that regulates the selective passage of substances while maintaining tissue integrity, which is essential for proper physiological function and protection against pathogens [2]. This barrier is maintained by an intricate network of junctional complexes, including AJs and TJs, that ensure cell-cell adhesion and control paracellular permeability [37]. The AJ is also linked to the ACTIN cytoskeleton *via* complexes involving proteins such as CATENIN ALPHA-1 and BETA-CATENIN [34]. Importantly, epithelial tissues undergo constant remodelling to maintain integrity in the face of dynamic environmental conditions and cellular stressors. This remodelling involves the coordinated trafficking of junctional proteins and the reorganization of the cytoskeleton, both of which are essential for preserving tissue architecture and barrier function. In this current report, we have significantly expanded on our previous studies to provide a more comprehensive understanding of how BECLIN-1 loss impacts these interconnected processes associated with maintaining epithelial integrity (see Supplemental Figure S1B for a schematic integrating all these new data).

In addition to E-CADHERIN [11], we now show BECLIN-1 loss leads to significant mislocalization of OCCLUDIN, resulting in their accumulation in the cytoplasm rather than their proper localization at cell junctions. This disruption is likely to reflect a failure in the endocytic trafficking machinery, where internalized proteins are no longer efficiently routed through early endosomes for recycling or degradation [38]. We previously identified that BECLIN-1-deficient cells exhibit defective RAB5A^+ve^ early endosome maturation, impairing E-CADHERIN trafficking. Here, live-cell imaging studies in MEFs further revealed that BECLIN-1 loss results in cargoes like Dextran and Transferrin accumulating near the plasma membrane and failing to progress through the endosomal pathway, implying a defect downstream of the internalization process. Interestingly, a recent report showed BECLIN-1 knockdown results in OCCLUDIN levels increasing on membranes in colorectal cancer cells (and decreasing when autophagy is induced using Tat-Beclin1 peptide) [13]. Whilst this finding is at odds with our results, it is an inherently different experimental system employing 2D adherent cell cultures that preclude investigation of the natural crypt-villus architecture. Indeed, we do see the accumulation of OCCLUDIN on lateral membranes, but its correct location at the apicolateral junction is impaired, which cannot be as readily discerned in the 2D cell cultures used in that study. Moreover, the cells are of colonic origin and cancerous, hence epithelial remodelling and turnover may be quite different to in normal IECs. As such, these factors complicate direct comparison with our system, which focussed on normal (i.e. non-cancerous) small-intestine derived cells.

The impact of E-CADHERIN mislocalization is of particular importance due to its downstream consequences with potential impact on the cytoskeleton. It is known that E-CADHERIN localization at the apicolateral membrane involves multiple pathways, including sorting and recycling *via* RAB5A^+ve^ and RAB11^+ve^ endosomes, respectively, and/or direct transport from the Golgi which also involves a RAB11-dependent mechanism [38]. As BECLIN-1 loss leads to widespread alterations in the distribution of endosomal compartments, including RAB5A^+ve^ and RAB11^+ve^ endosomes, this likely accounts for the decreased apicolateral distribution of E-CADHERIN. Importantly, the process of “cadherin flow” can compensate for the loss of E-CADHERIN at AJs by re-distributing E-CADHERIN from the lateral membrane towards the apical junction in an F-actin-dependent manner, thereby helping to re-establish cell-cell contacts [38, 39]. The increased lateral localization of E-CADHERIN following BECLIN-1 loss suggests the activation of an alternate recycling pathway, potentially *via* RAB4-mediated rapid recycling, which bypasses the RAB11^+ve^ endosomal recycling pathway [38, 40]. Cadherin flow also relies on the association of E-CADHERIN with F-actin *via* CATENIN ALPHA-1 and BETA-CATENIN, which was preserved following BECLIN-1 loss [39]. However, despite a compensatory increase in lateral F-actin, which suggests that cadherin flow may be enacted, this was insufficient to fully restore E-CADHERIN levels at the AJs.

The connection between BECLIN-1 loss and E-CADHERIN mislocalization is further complicated when the process of E-CADHERIN turnover is considered. This normally involves its degradation in lysosomes following delivery by RAB7^+ve^ endosomes [41]. We previously noted that the number of RAB7^+ve^ endosomes is increased following BECLIN1 loss [11], potentially as a compensatory mechanism in response to the increased levels of mislocalized E-CADHERIN. However, in this current study we demonstrated that lysosomal proteolytic activity is also impaired, which would further compound the apparent accumulation of E-CADHERIN in the cytoplasm. Some reports have also demonstrated that lysosomal activity associated with autophagy can also contribute to cargo degradation *via* RAB7^+ve^ vesicles [32, 42], which potentially explains why ATG7-deficient organoids also showed mislocalization of E-CADHERIN. Interestingly, we previously showed that the RAB7^+ve^ vesicles were smaller in size following ATG7 loss [11] providing further evidence that this compartment is altered in ATG7^ΔIEC^ organoids. However, the more profound impact of BECLIN1 loss is likely due to the additional effects of defective endocytic trafficking.

The contractile tension generated at the cell junction, critical for transmitting mechanical forces between cells and preserving tissue integrity, relies on the interaction between F-actin and E-CADHERIN *via* the catenin complex composed of both CATENIN ALPHA-1 and BETA-CATENIN. Combined, these proteins contribute to the formation of the apical F-actin belt [34, 37]. Of relevance to our model system, disruption of this F-actin belt has been associated with epithelial damage in models of colitis and in IBD patients [43–45]. Notably, this belt was reduced in the BECLIN-1 knockout organoids, which most probably contributes to the loss of GI tissue architecture both in the organoids and BECLIN-1-deficient mice. Mechanistically, this loss of the F-actin belt is likely a consequence of E-CADHERIN loss at the AJs, which not only recruit CATENIN ALPHA-1 and BETA-CATENIN for F-actin attachment but also proteins such as Rho effectors required for F-actin formation [46, 47]. This is consistent with previous reports showing BECLIN-1 interacts with the Rho GTPase, RAC1 (Rac family small GTPase 1), and BECLIN-1 loss leads to cytoskeletal defects in macrophages [48]. The reduction of E-CADHERIN at AJs likely also accounts for the reduced levels of BETA-CATENIN at the apical membranes.

Our data suggests that BECLIN-1 is pivotal in epithelial tissue remodelling particularly in trafficking junctional proteins and reorganizing the cytoskeleton to ensure proper localization and degradation of key proteins like E-CADHERIN and OCCLUDIN (Supplemental Figure S1B). Loss of BECLIN-1 disrupts epithelial remodelling, leading to impaired junctional integrity, increased permeability, and epithelial barrier dysfunction. These wide-ranging defects align with the gross and rapid disruption of GI epithelial architecture observed in BECLIN-1-deficient mice. Hence, these findings highlight the importance of BECLIN-1 in epithelial homeostasis. While BECLIN-1 has not previously been implicated in diseases characterized by barrier defects, such as IBD, our findings raise the possibility of its involvement in such conditions, particularly those affecting the small bowel such as Crohn’s disease, given the severe enteritis observed following BECLIN-1 deletion. This highlights the need for further investigation into BECLIN-1’s role in epithelial homeostasis and disease.

## Methods

### Mice

Details of the *Becn1^wtIEC^*, *Becn1^ΔIEC^*and *Atg7^wtIEC^ Atg7^ΔIEC^* mice used for generation of organoids and MEFs were described previously [11]. These were housed at the La Trobe Animal Research and Teaching Facility (LARTF, La Trobe University, VIC, Australia) under Specific Pathogen Free conditions. All experiments performed were approved by the La Trobe University animal ethics committees (approvals AEC18024, AEC18036) in accordance with the *Australian code for the care and use of animals for scientific purposes*. We have complied with all relevant ethical regulations for animal use.

### Intestinal organoid culture

Organoids were established from crypt-enriched fractions from the duodenum of untreated exactly as described previously [11] . Organoids were maintained at 37°C, 5 % CO_2_, with media being replaced every 2-3 days. Passaging was performed every 7-10 days by mechanically dissociating organoids and re-seeding into fresh Cultrex at a 1:3 split. To induce *Becn1* and *Atg7* deletion, organoids were seeded into media containing 200 nM 4-HT (Sigma-Aldrich, H7904) for three days, and then maintained as per normal.

### Mouse embryonic fibroblast generation and culture

Mouse embryonic fibroblasts were generated from E13-E14.5 embryos derived from *Becn1^wtIEC^*, *Becn1^ΔIEC^* and *Atg7^wtIEC^ Atg7^ΔIEC^* mice and immortalized (at passage 2-4) with SV40 large T antigen, as described previously [49]. Cells were maintained in DME Kelso medium supplemented with 10% (v/v) fetal bovine serum, 250 mM L-asparagine and 50 mM 2-mercaptoethanol. Deletion of *Becn1* and *Atg7* was achieved by culturing cells in the presence of 500 nM 4-HT for 3 days, and then maintained as per normal. Protein deletion was confirmed by Western blot analysis.

### Western immunoblotting

Lysates of MEFs and organoids were prepared, electrophoresed and transferred to membranes for Western blotting as described previously [11]. Following blocking in 5 % (w/v) skim milk in PBS (2.7 mM KCl, 1.76 mM KH_2_PO_4_, 136.7 mM NaCl, 8.07 mM anhydrous Na_2_HPO_4_) for 1 hour at room temperature (RT) with agitation, membranes were probed overnight at 4°C with the following antibodies at the following dilutions, prepared in 1 % (w/v) skim milk in PBST (PBS + 0.05% (v/v) Tween-20): BECLIN-1, 1:500 (CST, 3495); ATG7, 1:500 (Sigma-Aldrich, A2856), SQSTM1/p62, 1:500 (CST, 5114); MAP1LC3B, 1:500 (Novus Biologicals, NB100-2220); E-CADHERIN, 1:600 (CST, 3195). LAMP1, 1:1000 (Abcam, ab24170); OCCLUDIN, 1:500 (Invitrogen, 33-1500); GAPDH, 1:5000 (Invitrogen, MA5-15738). Membranes were washed in PBST and probed for 1 hour at RT with the following antibodies, prepared in 1 % (w/v) skim milk in PBST: Donkey anti-Rabbit IgG, 1:10,000 (GE Healthcare, NA943V), Goat anti-Mouse IgG, 1:10,000 (Sigma-Aldrich, A0168). Luminescent signals were visualised using the Western Lightning Plus-ECL (PerkinElmer) kit and ChemiDoc Imaging System (Bio-Rad). Images were processed using Image Lab (Bio-Rad).

### Intestinal organoid whole-mount immunofluorescence

The whole-mount intestinal organoid staining protocol is as described previously which is an adaption of the methods from Dekkers, Alieva [50] with modifications to clearing and mounting steps. Briefly, organoids were removed from matrix by incubation with ice-cold Gentle Cell Dissociation Reagent (STEMCELL Technologies,100-0485) with gentle rocking at 4°C for 60 minutes. Organoids were then fixed in 4% (w/v) paraformaldehyde solution (ProSciTech, C004) and blocked with Organoid Wash Buffer (DPBS (Gibco^TM^, 14190144) + 0.1% (w/v) TritonX-100 (Merck) + 0.2% (w/v) Bovine Serum Albumin (Merck, A3059)), followed by overnight incubation at 4 °C with gentle rocking with the following primary antibodies at the indicated dilutions: E-CADHERIN, 1:400 (Invitrogen, 13-1900); OCCLUDIN, 1:100 (Invitrogen, 33-1500); F-actin, using 1X Phalloidin-iFluor 647 solution (Abcam, ab176759); BETA-CATENIN, 1:100 (Proteintech, 51067-2-AP). Following primary antibody incubation, organoids were washed extensively and incubated overnight at 4 °C with gentle rocking using the following secondary antibodies: Goat anti-Rat IgG (H+L) cross-adsorbed secondary antibody, Alexa Fluor^TM^ 568 (1:400, Invitrogen, A-11077); Goat anti-Mouse IgG (H+L) cross-adsorbed secondary antibody, Alexa Fluor^TM^ 488 (1:400, Invitrogen, A-11004); Goat anti-Rabbit IgG (H+L) Cross-Adsorbed Secondary Antibody, Alexa Fluor^TM^ 488 (1:400, Invitrogen, A-11008) and nuclear stain DAPI (1 μg/ml, Merck, D9542). Organoids were then subjected to another extensive washing step prior to sample clearing and mounting steps. Images were then acquired using Zeiss LSM 980 with Airyscan 2 confocal microscope. FastAiryscan 2 SR-4Y imaging mode was utilized, and parameters such as laser power and gain were kept consistent amongst samples. Images were taken using the 40x (water) objective with 1.7x Zoom and imaged once to avoid excessive photobleaching. Z-stacks were acquired for each image at 5 µm intervals.

### Dynamic endocytic trafficking analysis

Mouse embryonic fibroblasts (MEFs) were seeded overnight in MEF culture media at 2000 cells/well in Nunc^TM^ Lab-Tek^TM^ II Chambered Coverglass slides (Thermo Scientific, 155409PK). To visualise the different trafficking pathways, the media was removed and the cells were washed with Live Cell Imaging Solution (LCIS) (Invitrogen, A59688DJ). The media was then replaced with LCIS containing either pHRodo-Dextran (for bulk fluid-phase endocytosis, 60 µg/ml, Invitrogen, P35368) or pHRodo-Transferrin (for receptor-mediated endocytosis, 60 µg/ml , Invitrogen, P35376 ) with WGA conjugated to AlexaFluor^TM^ 594 (for clathrin-dependent and independent pathways pathway, 2 µg/ml, Invitrogen, W11262), along with Hoechst 33342 (2 µg/ml, ThermoFisher, 62249) to stain nucleic acids. After 30 minutes incubation at 37°C, cells were imaged using a Zeiss LSM 980 confocal microscope under controlled conditions (37°C, 10% CO_2_), with z-stacks acquired at 2 µm intervals.

### Lysosome function studies

Intestinal organoids were cultured as described above. At day 5 post-4-HT treatment, organoids were carefully removed from Cultrex by performing a series of washes with ice-cold Advanced DMEM/F12 medium (Gibco, 12634010). The organoids were then resuspended in a staining solution containing LCIS, LysoTracker Red^TM^ DND-99 (50 nM, Invitrogen^TM^, L7528), DQ^TM^ Green BSA (50 µg/ml Invitrogen^TM^, D12050), and Hoechst 33342 (5 µg/ml) and transferred to a Nunc^TM^ Lab-Tek^TM^ II Chambered Coverglass. The organoids were incubated in the staining solution for 1 hour at 37°C in 5% CO_2._ Following incubation, images were acquired using a Zeiss 980 Confocal Microscope at 40x magnification, with matched pinhole sizes for all lasers and z-stacks captured at 4 µm intervals.

### Quantitative image analysis

Fluorescent image analysis was performed using ImageJ Software (Fiji). Spatial calibration was conducted by setting the appropriate scale for each image (Analyse > Set Scale). Quantification of apical, lateral, basal and cytoplasmic fluorescence signals on whole-mount fluorescent images (Figure 1A, Figure 3A, I) was conducted by manually drawing regions of interest (ROIs) around the relevant structures using the polygonal, freehand and line selection tools and measured using the ‘Measure’ (Analyse > Measure) function. A minimum of 3 ROIs (comprising of >5 cells one after another) per stack, and at least 3 z-sections with clear apical-to-basal orientation, were analyzed per organoid.

For endocytic uptake (Supplemental Figure S1C, F), z-stacks were compressed into a single layer using the Z-projection function (Image > Stacks > Z Project). ROIs were manually drawn around individual cells, with at least 10 cells analyzed per replicate. The fluorescence channel of interest was thresholded to generate a binary image (Image > Adjust > Threshold), followed by particle separation via watershed segmentation (Process > Binary > Watershed). Particles within the ROI were quantified using the Analyze Particle tool (Analyze>Analyze Particle).

For lysosome function analysis (Figure 2A), fluorescent signals from each z-section were thresholded to isolate structures of interest (i.e. LysoTracker Red DND-99^+ve^ puncta or DQ Green BSA^+ve^ puncta). Watershed segmentation (Process > Binary > Watershed) was performed next to separate particles. These particles were then quantified using the 3D Object Counter tool (Analyze > 3D Objects Counter) to determine the number and the size of the particles within the organoid z-stack. The number of particles was normalized to the organoid volume.

Co-localization analysis was performed using the Just Another Colocalisation Plugin (JaCoP; Bioimaging and Optics Platform, BIOP) with Otsu thresholding applied to define object boundaries.

### Statistics and reproducibility

Numerical source data for all graphs are provided in Methods and Figure legends. Statistical tests were performed using GraphPad Prism 8 Software (GraphPad, SanDiego, CA) via Student’s unpaired t tests between two groups and one-way ANOVA (with Tukey post-hoc comparisons) for multiple comparisons. All data were obtained by performing at least n = 3 biological replicates with representative data shown and expressed as the mean ± standard error of the mean (S.E.M). P values < 0.05 were considered statistically significant. Significance levels were split further as: **P < 0.01, ***P < 0.001, ****P < 0.0001.

## Acknowledgements

We acknowledge scholarship support for J.J. (La Trobe Graduate Research Scholarship and Full Fee Research Scholarship) and S.T. (La Trobe University Research Training Program Scholarship). We are grateful to the Australian Research Council for grant support (E.F.L. and W.D.F., DP190102612; P.A.G., DP160102394), the National Health and Medical Research Council (K.D., A.S.Y., 2010704; PDC 2008909), the US DOD (HT94252310088), and the Victorian Cancer Agency (E.F.L., MCRF19045) for fellowship support.

## Abbreviations

AJs: adherens junction
GI: gastrointestinal
IBD: inflammatory bowel disease
IECs: intestinal epithelial cells
MEFs: mouse embryonic fibroblasts
PtdIns3K: phosphatidylinositol 3-kinase
ROI: region of interest
RT: room temperature
TJs: tight junctions
WGA: wheat germ agglutinin
4-HT: 4-hydroxytamoxifen.

