## Supplementary Figures 1 & 2 for "BECLIN-1 is Essential for the Maintenance of Gastrointestinal Epithelial Integrity by Regulating Endocytic Trafficking, F-actin Organization and Lysosomal Function"

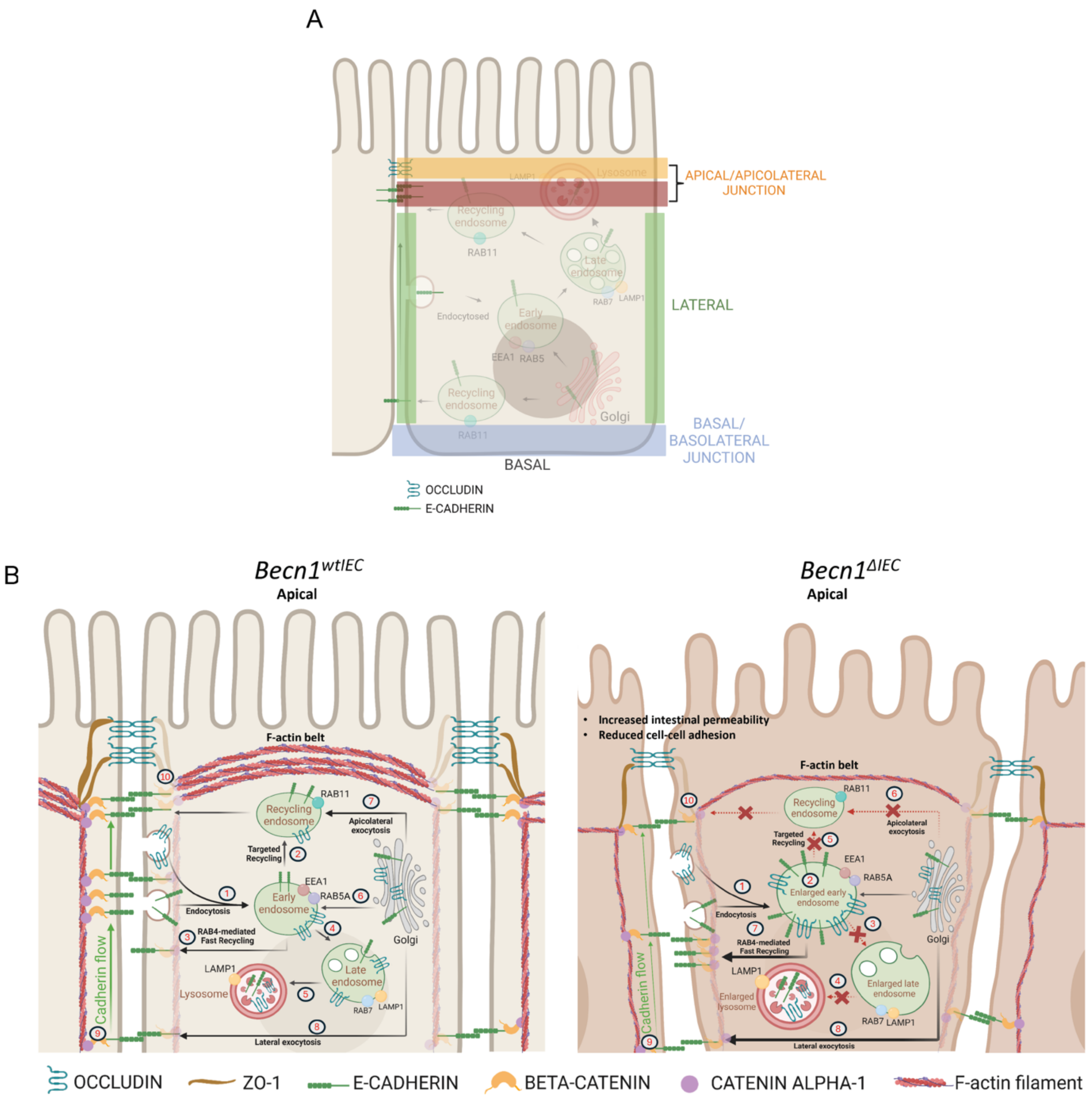


**Supplemental Figure S1.**

(**A**) Schematic illustrating the various membranes referred to in the main text. (**B**) Schematic representing the BECLIN-1 loss impacts endocytosis, formation of cellular junctions and the cytoskeleton. **Left:** In healthy IECs undergoing epithelial remodeling, (1) membrane-bound E-CADHERIN and OCCLUDIN are internalised *via* endocytosis and transported to RAB5A$^{+ve}$ early endosomes, where they undergo cargo sorting. The majority of E-CADHERIN and a portion of OCCLUDIN are (2) recycled back to the apicolateral junction through a RAB11-dependent targeted recycling pathway or (3) undergo rapid recycling to the lateral membrane *via* a RAB4-dependent pathway. (4) A large proportion of OCCLUDIN and some E-CADHERIN are sorted into RAB7$^{+ve}$ late endosomes for (5) transport to the lysosomes, where they are degraded. (6) The balance of cargo recycling and degradation is complemented by newly synthesized E-CADHERIN and OCCLUDIN from the Golgi, which is directed to RAB5A$^{+ve}$ early endosomes for sorting. Additionally, newly synthesized E-CADHERIN and OCCLUDIN can be (7) delivered to the apicolateral junction through RAB11-mediated apicolateral exocytosis or (8) delivered to the lateral membrane *via* lateral exocytosis. (9) Lateral E-CADHERIN can also be transported to the apicolateral junction through an F-actin-dependent cadherin flow mechanism. (10)

Once localized at the apicolateral junction, E-CADHERIN recruits F-actin polymerization machinery to form the cortical F-actin belt, generating the contractile tension essential for epithelial integrity. **Right:** Upon BECLIN-1 loss, (1) internalized E-CADHERIN and OCCLUDIN are still transported to RAB5A$^{+ve}$ early endosomes. However, due to defective endosomal maturation (RAB5A to RAB7 transition failure), (2) cargo accumulates in enlarged RAB5A$^{+ve}$ early endosomes, (3) preventing further transport to RAB7$^{+ve}$ late endosomes and (4) degradation in lysosomes. The late endosomes and lysosomes were also enlarged following BECLIN-1 loss. (5) Additionally, mislocalization of RAB11 in BECLIN-1-deficient IECs disrupts cargo sorting from RAB5A$^{+ve}$ early endosomes to RAB11$^{+ve}$ recycling endosomes, halting targeted recycling of cargoes to the apicolateral junction. (6) Newly synthesized E-CADHERIN and OCCLUDIN are also unable to undergo RAB11-mediated apicolateral exocytosis. As a compensatory mechanism, (7) lateral transport *via* RAB4 and (8) lateral exocytosis may be upregulated in BECLIN-1-deficient IECs. However, with reduced F-actin volume on the lateral membrane, (9) the cadherin flow mechanism to relocate cargo to the apicolateral junction becomes less effective. Together, these trafficking defects lead to a significant reduction in E-CADHERIN and OCCLUDIN at the apicolateral junction, impairing F-actin polymerization and hindering the formation of the cortical F-actin belt, contributing to epithelial breakdown. Red dashed arrows with crosses denote defective or inefficient trafficking, while thicker black arrows indicate increased trafficking.

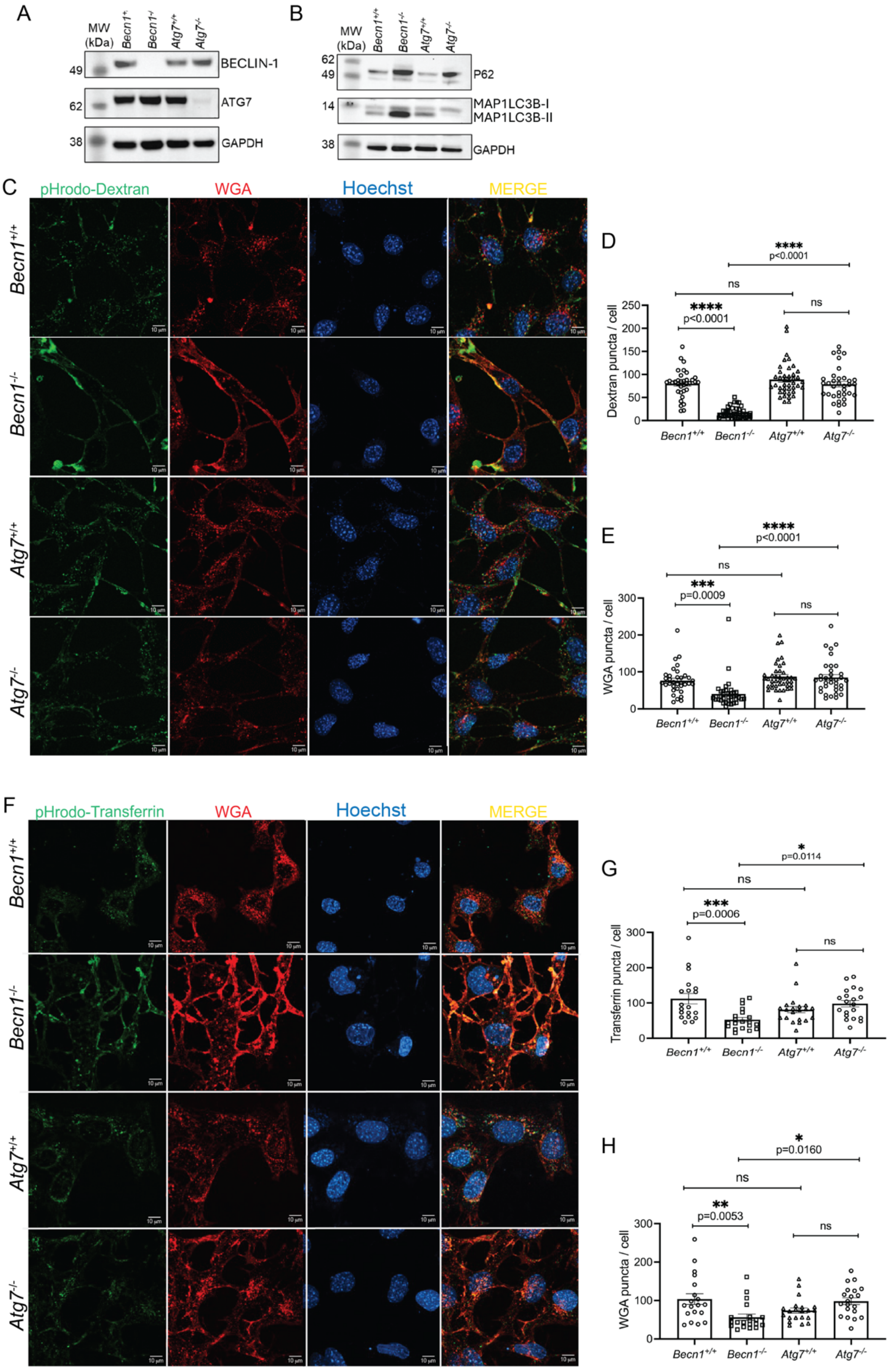


**Supplemental Figure S2. BECLIN-1 loss results in autophagy and endocytic trafficking defects in MEFs.**

(**A**) Deletion of BECLIN-1 and ATG7 at Day 5 post-4HT in MEFs, determined by Western blotting. This led to (**B**) defective autophagy, indicated by increased levels of total P62 and MAP1LC3, or increased MAP1LC3-I relative to MAP1LC3-II. GAPDH was used as a loading control.

Molecular weight (MW) markers indicate the relative size of the detected proteins. **C)** BECLIN-1 loss also led to defective trafficking of bulk fluid endocytosis of Dextran, which accumulated near the plasma membrane, **D)** with fewer puncta in the cytoplasm **E, H)** as was also the case with wheat germ Agglutinin (WGA). **F, G)** Similar results were observed for receptor-mediated endocytosis of Transferrin. Data were representative of at least *n* = 3 biological replicates. Graphs showed the mean ± S.E.M. Significance was determined by ordinary one-way ANOVA for all comparisons. Scale bar = 10 µm.
